# Analysis of *in-patient* evolution of *Escherichia coli* reveals potential links to relapse of bone and joint infections

**DOI:** 10.1101/2023.05.10.540183

**Authors:** Stanislas Thiriet-Rupert, Jérôme Josse, David Perez-Pascual, Jason Tasse, Camille Andre, Léila Abad, David Lebeaux, Jean-Marc Ghigo, Frédéric Laurent, Christophe Beloin

## Abstract

Bone and joint infections (BJIs) are difficult to treat and affect a growing number of patients, in which relapses are observed in 10-20% of the case. These relapses, which call for prolonged antibiotic treatment and increase the risk of emergence of resistance, may originate from ill understood adaptation of the pathogen to the host. Here, we studied three pairs of *Escherichia coli* strains corresponding to three cases of BJIs and their relapse to better understand in-patient adaptation.

Whole genome comparison presented evidence for positive selection with prevalence of non-synonymous and loss of function mutations. Phenotypic characterization showed that biofilm formation capacity was not modified, contrary to what is usually described in such relapse cases. Although virulence was not modified, we identified the loss of two virulence factors (namely an AFA afimbrial adhesin and a YadA-like adhesin) contributing to immune system evasion in one of the studied relapse strain. Other selected strategies likely helped the relapse strains to outcompete competitors through global growth optimization and colicin production. This work highlights the variety of strategies allowing in-patient adaptation in BJIs.

## Introduction

Bone and joint infections (BJIs), including prosthetic joint infections (PJI), are severe and difficult to treat infections affecting a growing number of patients, and showing 10 to 20% of relapse (1–6). Although staphylococci are responsible for more than 50% of all BJI, 20% of them are caused by *Enterobacterales*, especially in the case of early and late acute infections (7), whereas *Pseudomonas aeruginosa* infections are associated with particularly difficult to treat BJI (6). These data therefore support the need to study BJI caused by both Gram-positive and Gram-negative bacteria and for a better understanding of the *in vivo* adaptation leading to relapses.

During host colonization, pathogenic bacteria must face different selective stresses including host defenses and interactions with the endogenous microbiota or other co-infecting bacteria. This competition could involve toxin production (8, 9) and other adaptive strategies to access resources (10) and evade the host immune system (11, 12), eventually leading to a decrease virulence accounting for bacterial persistence in the host.

The genetic modifications underlying these adaptations are in general quite restricted upon natural host purifying selection (13, 14). In some extreme cases, mutator genotypes can evolve, thus facilitating within-host adaptation (15, 16). Extensive genome reductions and rearrangements (17–19) are also observed leading to the loss of non-essential genes in the host environment. However, there is an increasing body of evidence that these gene losses could also fuel adaptative evolution and microbial adaptation within patient is currently under intense scrutiny (20, 21). However, acquiring longitudinal samples and studying pathogens adaptation from the same patient in the context of infection relapses is difficult (22). In this study, we analysed three pairs of *E. coli* strains isolated from initial and recurrent BJIs in 3 patients to investigate how *in-patient* evolution occurred in these situations and provide insights on the potential mechanisms involved in infection relapse.

## Methods

### Selection of clinical strains

Initial / Relapse pairs of clinical *E. coli* strains used in this study were selected among the routine collections of the Bacteriology Department of Hôpital de la Croix-Rousse, Hospices Civils de Lyon and the Bacteriology laboratory at Cerballiance (Lyon, France). The first search criterion corresponded to collect *E. coli* strains isolated from BJI from 2017 to 2020. Then, initial/relapse pairs were formed when *E. coli* strain was isolated in two different samples at two different times in the same patient (initial and relapse) and at the same site of infection. Strain characteristics are available in Table 1. Antimicrobial susceptibility tests were performed by using the disc diffusion method (Supplementary Table 1).

**Table 1.** Cases and strain information. Table summarizing the type and location of infection as well as type of sample. Each strain was typed *in silico* and, when the infection was polymicrobial, the other species involved are listed.

| Case N° | Strain | Infection status | Infection site | Sample type | Diagnosis | Time before relapse | Antibiotic susceptibility profile | Phylogroup | Serotype | ANI | MLST #1 | MLST #2 | fimH | fumC | Infection type | Other pathogens if polymicrobial |
| --- | --- | --- | --- | --- | --- | --- | --- | --- | --- | --- | --- | --- | --- | --- | --- | --- |
| 1 | I1 | Initial | Hand | Deep tissue by swab | Septic arthritis 4th finger (no material) | 51 days | Penicillinase | B1 | O76:H28 | 99.99% | 224 | 730 | FimH3<br>9 | FumC4 | Monomicrobial |  |
|  | R1 | relapse |  | Deep tissue by swab |  |  | Identical |  |  |  |  |  |  |  | Polymicrobial | *Enterococcus faecium*,<br>*Pseudomonas aeruginosa*,<br>*Citrobacter freundii* |
| 2 | I2 | Initial | Spine | Washing liquid | Vertebral osteomyelitis (no material) | 748 days | ESBL-producing | G | O24:H4 | 99.99% | 117 | 48 | FimH9<br>7 | FumC4<br>5 | Monomicrobial |  |
|  | R2 | relapse |  | biopsy |  |  | Identical |  |  |  |  |  |  |  | Polymicrobial | *Cutibacterium acnes* |
| 3 | I3 | Initial | Right hip | Bone | Prosthetic joint infection | 66 days | ESBL-producing | A | O89/O16<br>2:H9 | 99.99% | 167 | 2 | / | FumC1<br>1 | Monomicrobial |  |
|  | R3 | relapse |  | Deep tissue |  |  | Identical |  |  |  |  |  |  |  | Monomicrobial |  |

### Bacterial strains and growth conditions

Bacterial strains used in this study are listed in Supplementary Table 2. Bacteria were grown in Miller’s Lysogeny Broth (LB) (Corning) or M63B1 minimum medium supplemented with appropriate antibiotic and carbon source as specified. Liquid cultures were incubated at 37°C or 30°C with 180rpm shaking. Solid cultures were done on LB with 1.5% agar supplemented with the appropriate antibiotics. Chemicals and media were purchased from Sigma-Aldrich.

### Strain construction

Insertion of a Zeo-GFP and Zeo-mars genetic cassettes at the permissive lambda *att* attachment site as well as the deletion of *fliC* in I1 and R1 strains and the deletion of two adhesin coding genes in I2 strain were done using pKOBApra plasmid (23) and lambda-red recombination. Primers used for genetic constructions are listed in Supplementary Table 3.

### RDAR morphotypes on congo red plates

Two microliters of an overnight culture grown at 37°C in LB medium under shaking (with appropriate antibiotics when needed) was spotted onto LB plates (without NaCl) containing 0.004% Congo Red and 0.002% brilliant blue. The spotted drops were allowed to dry, and the plates were incubated for 24 and 48 hours at 30 or 37°C.

### Auto-aggregation curves

Cultures were grown as described above, and 1mL aliquots were diluted to an OD_600_=3 in spent medium in order to prevent growth during experiment. Once diluted, all the tubes were vortexed and an aliquot of 50µL was removed, 1cm below the top of the culture, and diluted with 50µL of LB prior to OD_600_ measurement. The bacteria were then left to aggregate at RT for 6 hours, while repeating the measurement of OD_600_ 1 cm below the top of the culture every hour.

### Biofilm microtiter plate CV assay

Bacteria were grown overnight and diluted to an OD_600_=0.05 and 100µL were inoculated in technical triplicates in either a polystyrene Greiner 96-well plate or a polyvinyl chloride (PVC) round bottom 96-well plates (Corning). For each experiment a second identical plate was inoculated to assess growth. After 24 hours at 37°C, one plate was resuspended to measure the OD_600_. In the other plate, non-attached bacteria were washed away by flicking and wells were washed twice with water. Biofilms were stained with 125µL of crystal violet 1% (V5265; Sigma-Aldrich) for 15 minutes. Crystal violet was then removed by flicking and washed three times with water and plates were allowed to dry up. Biofilms were then resuspended using a solution of acetone:ethanol 1:5 mix, and OD_575_ was measured using Tecan Infinite-M200-Pro spectrophotometer. To account for potential differences in growth capacities, the OD_575_ values were normalized by OD_600_.

### CFU counting on biomaterials

The biofilm forming capacity was assessed on different orthopaedic biomaterials [stainless steel (AISI 316L), titanium (TA6V), medical grade silicone and ultra-high molecular weight polyethylene (UHMWPE)]. Uniform cylindrical pegs of orthopaedic biomaterial were manufactured by a supplier of medical and surgical orthopaedics equipment (Groupe Lépine, Genay, France) under the same conditions of manufacture of orthopaedic biomaterial and sterilized by gamma rays. Biomaterial pegs were fixed into the lid of 24-well flat-bottom cell culture plates (Corning Inc., Corning, NY, USA). Bacterial biofilms were formed by immersing the lid-containing pegs into 2mL of cultures at OD_600_=0.05 and incubated in LB for 24 hours at 37°C in humid atmosphere. After 3 rinsing steps in PBS, the pegs were dropped in a Falcon tube containing 5mL of PBS for biofilm disruption using sonication for 5 minutes (BACTOSONIC 14.2, BANDELIN, Berlin, Germany). Then, viable cell count was determined by serial dilution and plating.

### Bacterial adhesion to MG63 human osteoblastic cells

The osteoblastic cell line MG63 (LGC standards, Molsheim, France) was used to test bacterial adhesion. Cells were cultured in 75cm² flasks (T75, BD Falcon, Le Pont de Claix, France) at 37°C under 5% CO_2_, in a culture medium composed of DMEM (Dulbecco’s Modified Eagle Medium containing D-glucose, L-glutamine, pyruvate) supplemented with 10% fetal calf serum (FCS), penicillin (100µg/mL) and streptomycin (100µg/mL), all from Gibco (Paisley, UK). Cells were passaged once a week and used until passage 25 at the most. The day before the infection, MG63 cells were seeded at 100,000 cells per well in a 24-well plate and the different bacterial strains were inoculated in LB. The day of the infection, overnight bacterial cultures were centrifugated, resuspended and adjusted in DMEM + 10% FCS to reach a MOI (multiplicity of infection) of 100:1 and added to cell culture. After 2 hours of infection, cells were washed twice with PBS and incubated and lysed by osmotic shock in sterile water and lysates were plated on tryptone soy agar plates (Biomérieux) using the Easy Spiral® automaton (Interscience, Saint-Nom-la-Bretèche, France). CFU were counted after 18 hours of incubation at 37°C and results were presented as CFU for 100,000 cells.

### Growth capacity

An overnight culture of each strain was diluted to OD_600_ of 0.05 in appropriate medium. One hundred and fifty μL aliquots were inoculated in a 96-well plate. The plates were then incubated in a TECAN Infinite M200 Pro spectrophotometer (Männedorf, Switzerland) for 18 hours at 37°C with shaking of 2mm amplitude. The absorbance of each culture at 600 nm was measured every 15 minutes. Growth curve of each strain was measured using 3 biological replicates. The R package GrowthCurver (24) was used to infer growth-related parameters for all strains based on the growth curves.

### Cytotoxicity on HeLa cells

The cytotoxic activity on Hela cells of each bacterial strain was analyzed using the CellTiter-Blue® Cell Viability Assay (Promega) following manufacturer instructions. HeLa cells were cultured in complete DMEM (Life Technologies) supplemented with 10% FCS, 100µg/mL penicillin/streptomycin, and 10mM L-glutamine at 37C in the presence of 5% CO_2_. For infection assay, the HeLa cells were incubated in 96-well plates to reach a density of 10^5^ cells. Each bacterial strain was grown overnight in LB as described above and then washed twice in PBS, to finally adjust the bacterial density to 10^7^ CFU/ml. These bacterial suspensions were used to infect HeLa cells by adding 100 µl of each to the wells (MOI of 1:100). The plates were incubated at 37C in the presence of 5% CO_2_ for 2 hours. Non-infected cells and the *E. coli* K12 MG1655 strain were used as negative controls. At the end of the incubation time, the wells were washed twice with PBS to remove the bacteria. Then, 100 µl of PBS and 20 µl of CellTiter Blue solution were added to each well and incubated for 2 hours at 37C and 5% CO_2_. The cell viability was measured by recording the fluorescence at 560/590 nm using a Tecan Infinite-M200-Pro spectrophotometer.

### Hemolysis capacity

The hemolytic activity of each strain was assessed by mixing defibrinated horse blood (bioMerieux, Marcy l’Etoile, France) diluted to 10% (v/v) in PBS with a total of 10^8^ bacteria in PBS. After 2, 6 and 24 hours of incubation at 37°C with shacking, the non-lysed erythrocytes were pelleted by centrifugation at 1500g for 2 minutes, and the OD_450_ of 150μL of the cell-free supernatants was measured. The hemolytic capacity of each adapted strain was expressed relatively to its ancestral strain.

### Bacterial survival in human serum

For each strain, 50µL at OD_600_=0.4 was mixed in a well of a 96-well plat with 50µL of active human serum. The same process was followed for a control with heat-inactivated serum (HIS) (30 minutes at 55°C). The plate was then incubated at 37°C under shaking (450rpm) for 1h30. The content of each well was then serially diluted and plated on LB for CFU count and the survival of each strain was expressed as the ratio between the CFUs in active serum divided by the CFUs in HIS.

### Zebrafish care and maintenance

Homozygous Tg(*mfap4::mCherryF*) (ump6Tg) (25) Tg(*mpx::GFP*)^i114^ (26) double transgenic fishes were raised in Institut Pasteur zebrafish facility. Eggs were obtained by natural spawning, bleached according to standard protocols, and then kept in Petri dishes containing Volvic spring water and, from 24 hpf onwards, 0.003% 1-phenyl-2-thiourea (PTU) (Sigma-Aldrich) was added to prevent pigmentation. Embryos as well as infected larvae were reared at 28°C. Larvae were anesthetized with 200 μg/mL of buffered tricaine methane sulfonate (MS-222, Sigma-Aldrich) during the injection procedure as well as during *in vivo* imaging and processing for bacterial burden evaluation.

### Zebrafish infection

Overnight bacterial cultures were centrifuged, washed, and resuspended at the desired concentration in PBS. Anesthetized zebrafish larvae were microinjected intravenously 52–58 hours post-fertilization with 1nL of each bacterial suspension at a final concentration of 2−10^3^ CFU (27). Infected larvae were transferred into individual wells containing 1mL of mineral Volvic water (Danone, France) and 0.003% 1-phenyl 2-thiourea (used to improve optical transparency in zebrafish) in 24-well plates, incubated at 28°C, and regularly observed under a stereomicroscope. Protocols describing zebrafish maintenance and analysis, see the corresponding supplementary method section.

### Evaluation of the bacterial burden in infected zebrafish larvae

Infected zebrafish larvae were collected at 0, 24, and 48 hpi and lysed for the evaluation of the bacterial burden as previously described (28). Each larva was placed in an individual 1.5 mL Eppendorf tube and anesthetized with tricaine (200 μg/mL), washed with 1 mL of sterile water, and placed in 150 µL of sterile water. Larvae were then homogenized using a pestle motor mixer (Argos). Each sample was transferred to an individual well of a 96-well plate and 10 X serial dilutions were performed. For CFU enumeration and to assess plasmid stability throughout the infection kinetics, serial dilutions of lysates were plated on MacConkey agar plates supplemented with lactose plates and incubated overnight at 37°C.

### Visualization of macrophages and neutrophils in infected zebrafish larvae

Visualization of macrophages and neutrophils on living transgenic reporter larvae was performed upon infection as we previously described (29). Briefly, bright field, GFP, and RFP images of whole living anesthetized larvae were taken using a Leica Macrofluo Z16 APOA (zoom 16:1) macroscope equipped with a Leica PlanApo 2.0 X lens, and a Photometrics CoolSNAP *HQ2* camera. Images were captured using the Metavue software version 7.5.6.0 (MDS Analytical Technologies). After capture of images, larvae were washed and transferred to a new 24-well plate filled with 1mL of fresh water per well and incubated at 28 °C.

### Competition assay for biofilm formation

The pair of strains used in competition assay carried two different fluorescent tags and were inoculated in LB medium supplemented with appropriate antibiotic and incubated overnight at 37°C under shaking. Cultures were then adjusted to OD_600_ = 0.05 and mixed in 1:1 ratio (verified by FACS). Each competition mix was then inoculated in a PVC plate and incubated at 37°C for 16 hours. The supernatant was removed from each well, the biofilm resuspended in 100µL of PBS and the proportion of each strain was assessed by FACS and the relative fitness calculated as follows:

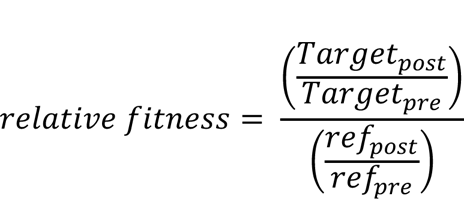

, where *Target_pre_* and *Target_post_* are the cell concentrations of the target strain in the mixed culture before and after the overnight incubation, and *ref_pre_* and *ref_post_* are the cell concentrations of the corresponding reference strain. In order to account for growth differences due to the expression of one or the other fluorescent tag (Supplementary Figure 1), competitions were run with both possible combinations of fluorescent tags. This protocol was used to compete R1 and R3 strains against their respective ancestors. However, because GFP and RFP tags were not expressed in I2 and R2 strains, we used an alternative strategy based on PCR amplification and Sanger sequencing of an SNP locus allowing to distinguish both strains (see Supplementary method section).

### Competition assay for growth in liquid cultures

The strains were incubated at 37°C in the appropriate medium and adjusted to OD_600_=0.05. A 1:1 ratio mixed culture was verified by FACS and incubated overnight at 37°C with shaking. The concentration of target and reference strains was measured using FACS and relative fitness was calculated as above.

### Alternative strategy for competition assays of I2 vs R2 strains

Even if we managed to introduce GFP and RFP (mars) tags in I2 and R2 strains, the related proteins were likely not expressed since thefluorescence was impossible to detect. Consequently we used an alternative strategy to assess the proportion of each strain. We took andvantage of the fact that both strains can be differentiated by SNPs. An aliquot of the mixed culture containing 10^7^ cells was pelleted and resuspended in 50µL of sterile distiled water and boiled for 10 minutes. Specific primers were then used to amplify a SNP locus in butC gene and Sanger sequence the resulting DNA fragment. Then, using QSVanalyzer (30), we calculated the frequeny of each mutation and used it to compute the relative fitness as above. To assess the accuracy of this strategy, we used strains I1 and R1 taged with different fluorescent marker to compare strains proportions as assessed by FACS and QSVanalyzer and found a very good correspondance between both methods (Supplemenatry Figure S6), therefore validating our alternative approach.

### Motility assay

Bacterial strains were incubated overnight in LB at the specified temperature with shaking. Then, 2µL of overnight culture was inoculated in swim plates (1% tryptone, 0.25% NaCl, 0.25% Eiken agar) for 24 hours at 37°C or 30°C. After incubation, motility was assessed by checking the spread of inoculated bacteria into the motility agar.

### Overlay assay colicin

I1 and R1 strains were incubated overnight at 37°C with shaking in LB supplemented with 40 µg/mL of Mitomycin C to induce colicin production. After incubation an overlay of each strain was realized by pouring 6mL of overnight culture diluted at OD_600_=0.1 onto a LB agar plate and removing it after 5 minutes. Plates were allowed to dry and a 10µL drop of supernatant from each strain culture filtered using 0.22uM filter was spotted on each overlay. After an additional drying step, the plates were incubated overnight at 37°C and the production of colicin was identified by the presence of an inhibition halo when the filtered supernatant was spotted. LB was spotted as a negative control and filtered supernatant from both strains was also spotted on overlay of each of them to be sur that the growth inhibition halo was not due to the presence of Mitomycin C in the filtered supernatant.

### Western blot analysis and LPS gel

The heat extracted proteins from 1mL at OD_600_=1 culture were suspended in 1×Laemmli buffer and incubated for 5 min at 95°C. The protein extracts were run on Mini-PROTEAN TGX Stain-Free precast Gels (BioRad) in 1×TGX buffer and then transferred to nitrocellulose membrane using a Trans-Blot^®^ Turbo Transfer System (BioRad). Blocking was performed in a 5% solution of dry milk and 0.05% Tween 1×PBS (1×PBST) overnight at 4°C with agitation. The membranes were then incubated in 1×PBST with either rabbit anti-flagellin (kindly given by Dr Monica Rolando) or mouse anti-RNAPα (BioLegend #663104) diluted 1/5,000 for 1h at room temperature with agitation. Membranes were washed in 1×PBST and then incubated with the secondary antibody (anti-rabbit or anti-mouse IgG conjugated with horse radish peroxidase at 1:10 000, Promega). The membranes were then revealed using an ECL kit (GE Healthcare) and the iBright^TM^ CL1500 system (Thermofisher).

LPS analysis on Mini-PROTEAN TGX Stain-Free precast Gels (BioRad) were realized as in (31).

### Whole genome sequencing and analysis

Prior to genome extraction, strains were inoculated in LB medium for over-day culture till the OD_600_ reached around 1.0 (ca. 5.0−10^8^ bacteria/mL). The bacterial cells were collected from 2mL of the culture and the genomic DNA was extracted using the Wizard Genomic DNA Purification Kit (Promega). Samples were then sent to Novogene Co., Ltd to be sequenced using Illumina NovaSeq 6000. Sequencing reads were pre-processed to remove low-quality or artefactual bases. We used fqCleaner v.0.5.0, a mini workflow implemented in Galaxy (32) (https://galaxy.pasteur.fr/) to process fastq files (quality trimming, duplicate and artifact filters). Draft genomes were de novo assembled using SPAdes version 3.15.0 (33) and annotated using Proka version 1.14.5 (34). Assembly quality and completeness was assessed using BUSCO version v5.4.2 (35). The assembled genomes were used for in silico typing and characterization of each strain using tools available at the Centre of Genomic Epidemiology (http://www.genomicepidemiology.org/): Resfinder (36–38), MLST (38, 39), SerotypeFinder (40), CHtyper (41). Phylogroups were identified in silico using ClermonTyping tool (42). In order to identify contigs corresponding to potential plasmids, each contig was used as a query in a BLASTn search using blast+ version 2.2.31 (38) against the COMPASS database (43). Only hits with 95% identity and 80% coverage of both the query and the hit sequence were kept.

Mutations in the adapted strain were identified by using the genome assembly of its ancestral strain as a reference. *breseq* version 0.35.0 (44) with the consensus mode was used with default parameters. The resulting identified mutations were filtered out if they were too close to each other (less than 50bp). These mutations are usually the result of misaligned reads, often due to repetitive regions (45). Because these regions are not 100% identical, they are disturbing the mapping process resulting in false positives. Mutations found in mobile elements were kept if only one copy was present in the genome.

### Statistical analysis

Figures and statistical tests were performed using either Prism 9.5.1 for Mac OS X (GraphPad Software, Inc.) or Rstudio with R version 4.2.1 using the R package GGplot2 (46).

### Data availability

All sequencing reads as well as genome assembly for each strain were deposited in NCBI under the BioProject accession number PRJNA932763. The code used in this study is available at https://github.com/Sthiriet-rupert/BJIs_adaptation.

## RESULTS

### I -*In-patient* evolved *Escherichia coli* clones showed no antibiotic resistance acquisition but signs of natural selection

In order to investigate potential cases of *in-patient* adaptation, we obtained three pairs of *E. coli* strains sampled from three patients presenting relapsing BJIs. Each pair is composed of a sample at the time of the initial infection (strains hereafter denominated as I1 to I3 for cases 1 to 3) and a second sample corresponding to the relapse of this infection (strains hereafter denominated as R1 to R3 for cases 1 to 3). The genome of each strain was sequenced and analyzed to confirm that the infection and relapse strains were indeed the same and not a case of re-infection (Table 1). Interestingly, antimicrobial susceptibility testing (Supplementary Table 1) and *in silico* analysis after whole genome sequencing (Supplementary Data 1) did not reveal any acquisition of antibiotic resistance associated with the relapse events. Consequently, each strain pair was examined for other phenotypic and genetic features that could have been selected to allow the infection relapses.

The first notable difference identified by genomic analysis was the number of gained and lost genes (Supplementary Table 4), some of which corresponding to potential plasmids (Supplementary Data 2). Particularly, the R2 strain that displayed the longest in-patient evolution underwent an important genome reduction with the loss of 3.4% of its size corresponding to 170 coding sequences involved in various functions such as transport, iron acquisition, carbohydrate acquisition and utilization, defense against competitors with Microcin H47 production-related genes and three adhesins (Supplementary Data 2).

The number of mutations per base per year was quite high for all three relapse strains (1.1−10^−6^ to 8.4×10^−6^) as compared to what was reported in the literature (6.9×10^−7^ for *E. coli* ED1a in the gut, (45)). In addition, even if no functional convergence was observed in the mutated genes, the predominance of loss of function and non-synonymous mutations suggests that positive selection was at play in these cases (Table 2).

**Table 2.** SNP analysis in the relapse strains. SNPs, insertions and deletions identified in the evolved strains as compared to their related ancestor.

| Strain | Mutation | Annotation | Impact | Gene | Description |
| --- | --- | --- | --- | --- | --- |
| R1 | T→A | intergenic (+168/-221) | Intergenic | uidR → / → uidA | transcriptional repressor/beta-D-glucuronidase |
|  | A→G | N16D (AAC→GAC) | Non-synonymous | yehT → | putative response regulator in two-component system with YehU |
|  | T→C | T46A (ACA→GCA) | Non-synonymous | rppH ← | RNA pyrophosphohydrolase |
|  | T→C | K577E (AAG→GAG) | Non-synonymous | ydiJ ← | putative FAD-linked oxidoreductase |
| R2 | G→A | G335G (GGC→GGT) | Synonymous | COMLMOEC 00122 ← | hypothetical protein |
|  | C→G | V16L (GTA→CTA) | Non-synonymous | insK 3 ← | IS150 transposase B |
|  | C→G | S67T (AGC→ACC) | Non-synonymous | COMLMOEC 04582 ← | IS3 family transposase IS629 |
|  | C→T | A131T (GCC→ACC) | Non-synonymous | ebgR ← | transcriptional repressor |
|  | C→T | A197V (GCC→GTC) | Non-synonymous | COMLMOEC 04752 → | IS66 family transposase ISEc47 |
|  | T→A | I263N (ATT→AAT) | Non-synonymous | btuC → | vitamin B12 ABC transporter permease |
|  | A→G | intergenic (+1283/-) | Intergenic | COMLMOEC 01687 → / - | hypothetical protein/- |
|  | A→C | intergenic (+352/-) | Intergenic | COMLMOEC 04947 → / - | hypothetical protein/- |
| R3 | G→T | E297D (GAG→GAT) | Non-synonymous | hypT → | hypochlorite-responsive transcription factor |
|  | G→A | D119N (GAT→AAT) | Non-synonymous | insF1 1 → | IS3 transposase B |
|  | +G | coding (1001/1041 nt) | Loss of function | vgrG1 → | Actin cross-linking toxin VgrG1 |
|  | C→T | intergenic (+315/-76) | Intergenic | uidR → / → uidA | transcriptional repressor/beta-D-glucuronidase |
|  | (GTGGCTGTT)8→10 | coding (1850/2208 nt) | Loss of function | traD ← | Coupling protein TraD |
|  | Δ1 bp | intergenic (-142/+229) | Intergenic | araD ← / ← araA | L-ribulose-5-phosphate 4-epimerase/L-arabinose isomerase |

### II -*In vivo* adaptation did not modify motility and biofilm formation capacities

Because strain motility could impact virulence, we tested each strain on low-agar LB plates at 30°C and 37°C, which showed that the second and third pairs of strains were non-motile (Supplementary Figure 2). Both I1 and R1 strains display temperature-dependent motility very likely due to a flagellin production restricted to 30°C (Supplementary Figures 2 and 3, see Supplementary Results).

Since biofilm formation is related to bacterial motility and sometimes involved in infection relapses, we analyzed the biofilm-forming capacity of each strain on different surfaces, which showed that the R1 relapse strain formed 1.71 more biofilms than I1 strain when grown on silicon coupons (Figure 1.A). No difference was identified for the I2-R2 pair (Figure 1.B), while the R3 relapse strain had a significant but very low increase in biofilm formation on PVC plates (1.08-fold) (Figure 1.C). This slight increase was reversed when grown on silicone since the I3 strain showed a 1.49 increase in biofilm formation (Figure 1.C). However, in all cases the tested clinical strains showed a weak biofilm-forming capacity, all of them being equivalently or even less efficient than *E. coli* K12 strain MG1655 used as a control for weak biofilm formation. Interestingly, R1 strain showed a slight decrease of its adhesion capacity to osteoblasts (Figure 1.D). However, no evident genetic modification could be linked to this phenotype. No difference was observed either in terms of auto-aggregation or production of curli amyloids and cellulose (Supplementary Figure 4 and Supplementary Results). Taken together, these results suggest that a modification of biofilm formation and biofilm-related phenotypes were unlikely to be involved in *in vivo* adaptation in these three studied cases.

**Figure 1.**
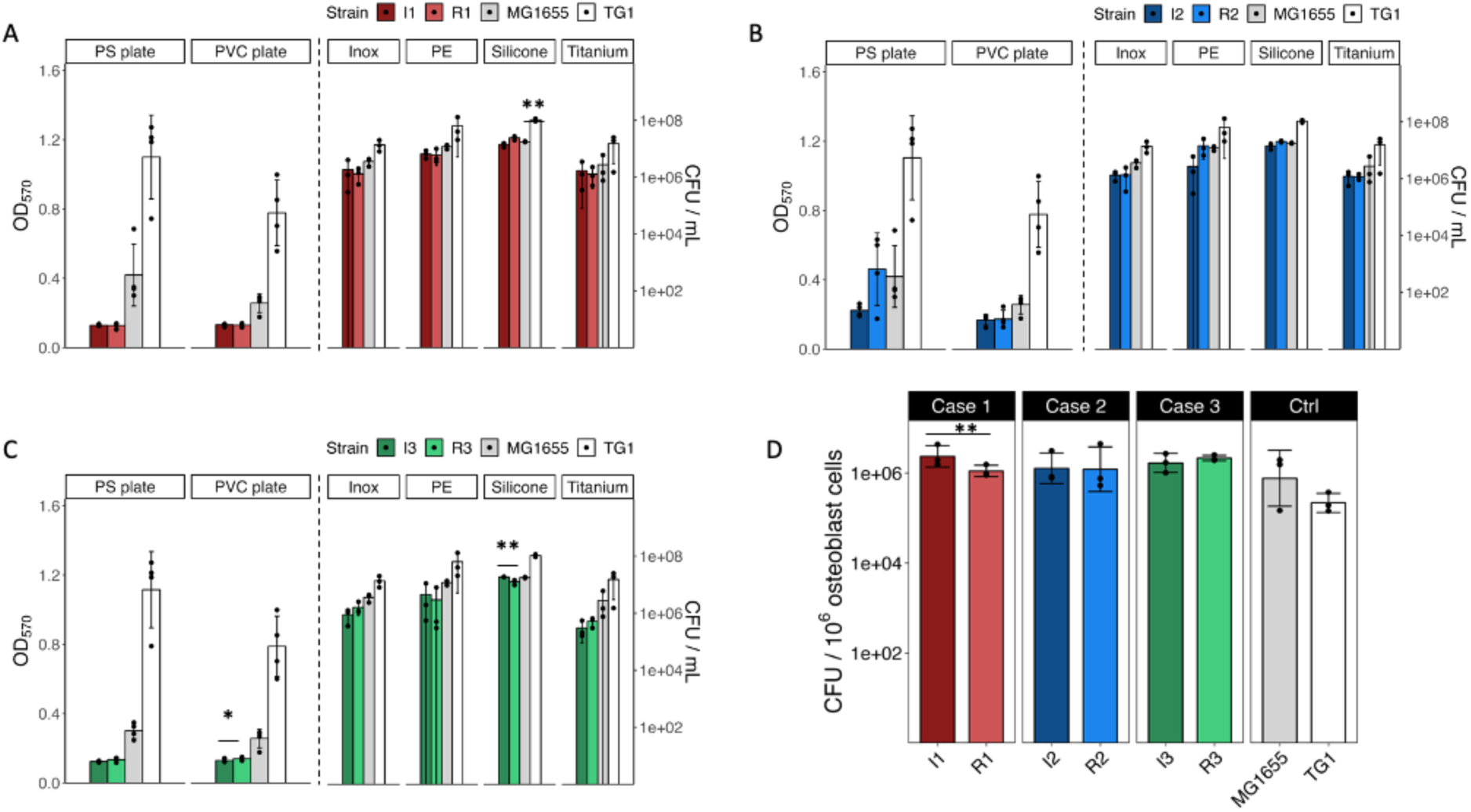
Biofilm formation capacity for infection and relapse strains from cases 1 (**A**), 2 (**B**) and 3 (**C**). Six different surfaces were tested, *E. coli* K12 MG1655 was used as a control for weak biofilm formation and *E. coli* TG1 as a control for strong biofilm formation. Biofilm formation on PS and PVC plates was assessed by crystal violet assay (n=4 biological replicates) and is plotted on left y-axis, while the biofilm formation on the other surfaces was assessed by CFU count (n=3 biological replicates) and are plotted on the right y-axis. (D) Adhesion capacity on osteoblast human cells (n=3 biological replicates). Statistical significance according to non-parametric Mann-Whitney test. * p<0.05; ** p<0.01.

### III -Loss of adhesins induces serum resistance in R2 strain

To assess the strain capacity to challenge to the host immune defenses, we measured the survival of all strains exposed to active human serum. While we observed no difference between I3 and R3 strains, the relapse strain R1 had a slight decrease in serum resistance compared to I1 strain and the relapse strain R2 had a clear increase in serum resistance relatively to its ancestral strain I2 (Figure 2.A). Although we identified no difference in the I2 *vs* R2 lipopolysaccharide profile (outer-membrane component targeted by the complement) (Figure 2.B), the loss of two adhesins in R2 could be linked to this phenotype (Supplementary Data 2) (11). Indeed, deletion of each adhesin in I2 strain led to reduced serum resistance suggesting that the loss of both adhesins could have been selected through the conferred immune system evasion, contributing to the strain adaptation to the host environment (Figure 2.C). Unfortunately, we were unable to assess the additive effect of these mutations since we could not reconstruct the double mutant in the I2 strain despite repeated attempts.

**Figure 2.**
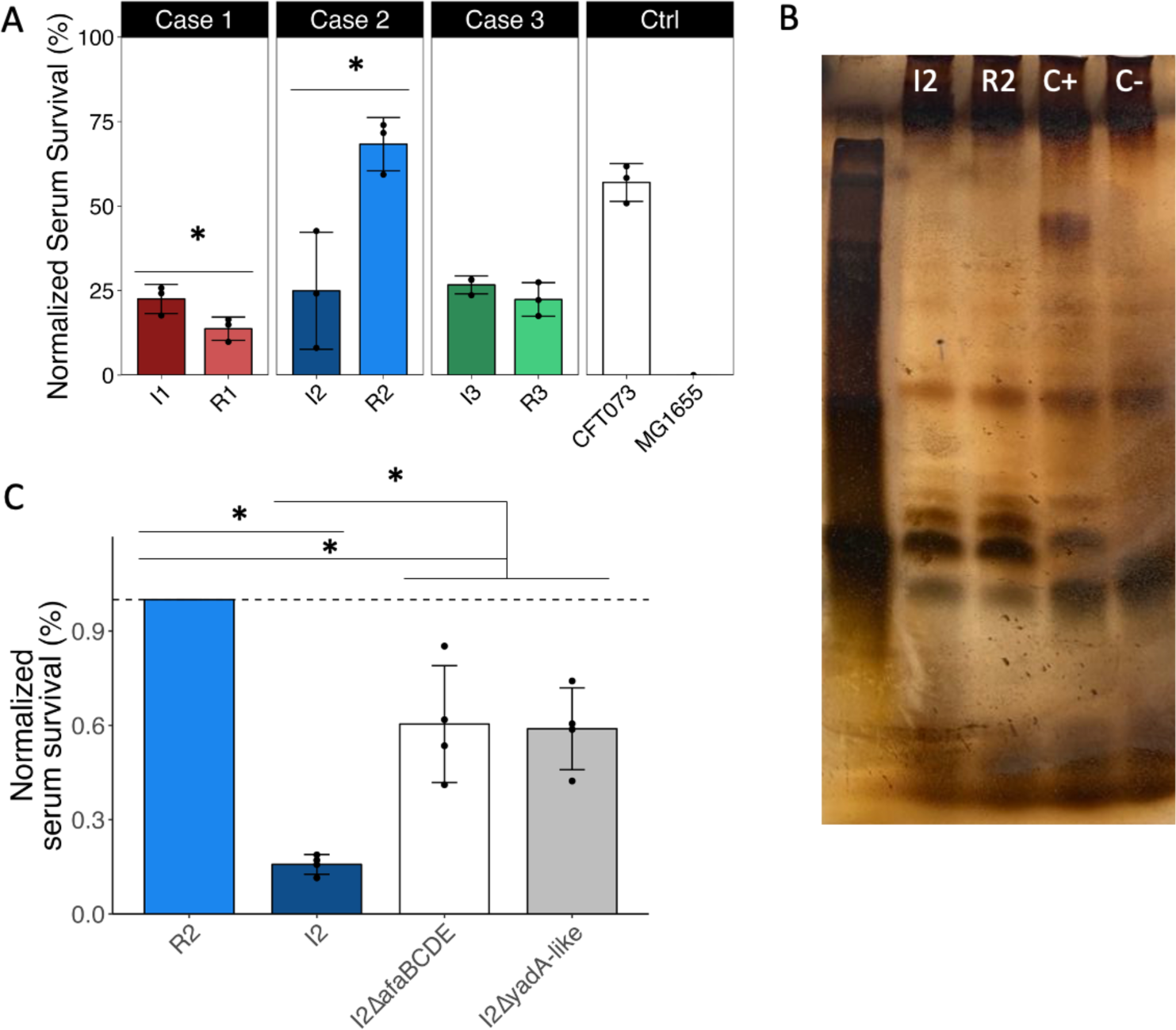
Serum survival capacity. **A**. Normalized serum survival capacity of each strain expressed as percentage of the control for growth in Heat Inactivated Serum (HIS). *E. coli* strains CFT073 and MG1655 were used as positive and negative controls, respectively. **B**. LPS profile of I2 and R2 strains. *E. coli* MG1655 was used as a negative control and the same strain in which the O-Antigen was reconstructed was used as a positive control. **C**. Serum survival of I2 ancestral strain, R2 relapse strain as well as two deletion mutants constructed in the I2 ancestral strain: one for the *afa* afimbrial operon and a second for the *yadA*-like encoding gene.

### IV -Strain virulence was not targeted during in-patient selection

To gain insight into the potential evolution of the virulence of the studied strains, we assessed their cytotoxicity on HeLa cells as well as hemolysis capacities and showed no differences, except for a slightly lower hemolysis induced by R2 strain after 24h of incubation in presence of defibrinated horse blood (Figure 3.A.B).

**Figure 3.**
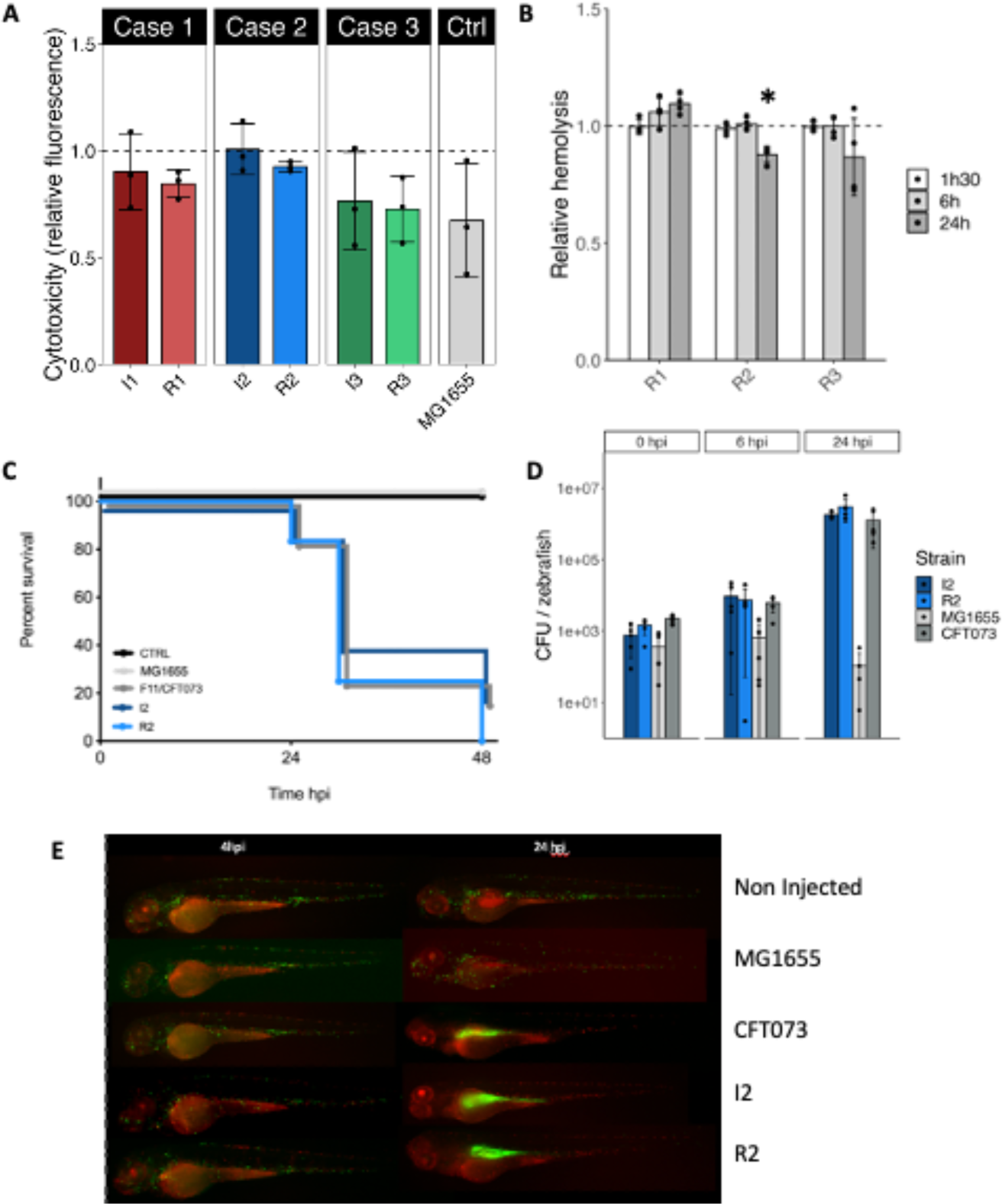
Virulence of the strains. (**A**) Cytotoxicity test on HeLa cells. All six adapted and ancestral strains were tested as well as MG1655 strain as a negative control. The values for each strain were expressed relatively to the control without bacteria. Values are represented as a mean of three biological replicates +/− SD. (**B**) Hemolysis capacity in liquid culture of each relapse strain relatively to its ancestor. Values are represented as a mean of four biological replicates +/− SD. (**C**) Monitoring of zebrafish survival, (**D**) bacterial burden into the zebrafish and (**E**) *in vivo* fluorescence of macrophages and neutrophils for zebrafish infected by I2 and R2 strains as well as non-injected control, MG1655 strain (negative control) and CFT073 (positive control). Each image is representative of 5 biological replicates. Fifteen zebrafish were used for each experiment in C and D.

Potential differences in *in vivo* virulence were tested in a zebrafish (*Danio rerio*) larval model (47). While no differences were found in bacterial load for all pairs (Sup Figure S5.A.B.C), zebrafish larvae exposed to R2 relapse strain had a slightly lower survival 24 hours post-infection (hpi) than with its related ancestor I2, even though both strains ultimately induced the same level of mortality 48 hpi (Figure 3.C.D). No difference was identified concerning the zebrafish innate immune response level (Figure 3.E). These results suggest that the R2 relapse strain induced slightly quicker mortality, which does not seem to be linked to a difference in colonization ability.

### V -Growth optimization and colicin production as strategies to outcompete other bacteria

To assess if a difference in growth capacity could advantage the strains isolated after relapses, they were pitted against their related ancestor in 1:1 ratio competition experiments in biofilm and liquid conditions. First, no difference in fitness was observed in biofilm condition, which further straightened the hypothesis that biofilm formation was not targeted during the *in vivo* adaptation. In liquid, while R1 strain slightly outcompeted its ancestor I1, R2 strain had a striking fitness advantage over I2 (Figure 4.A).

**Figure 4.**
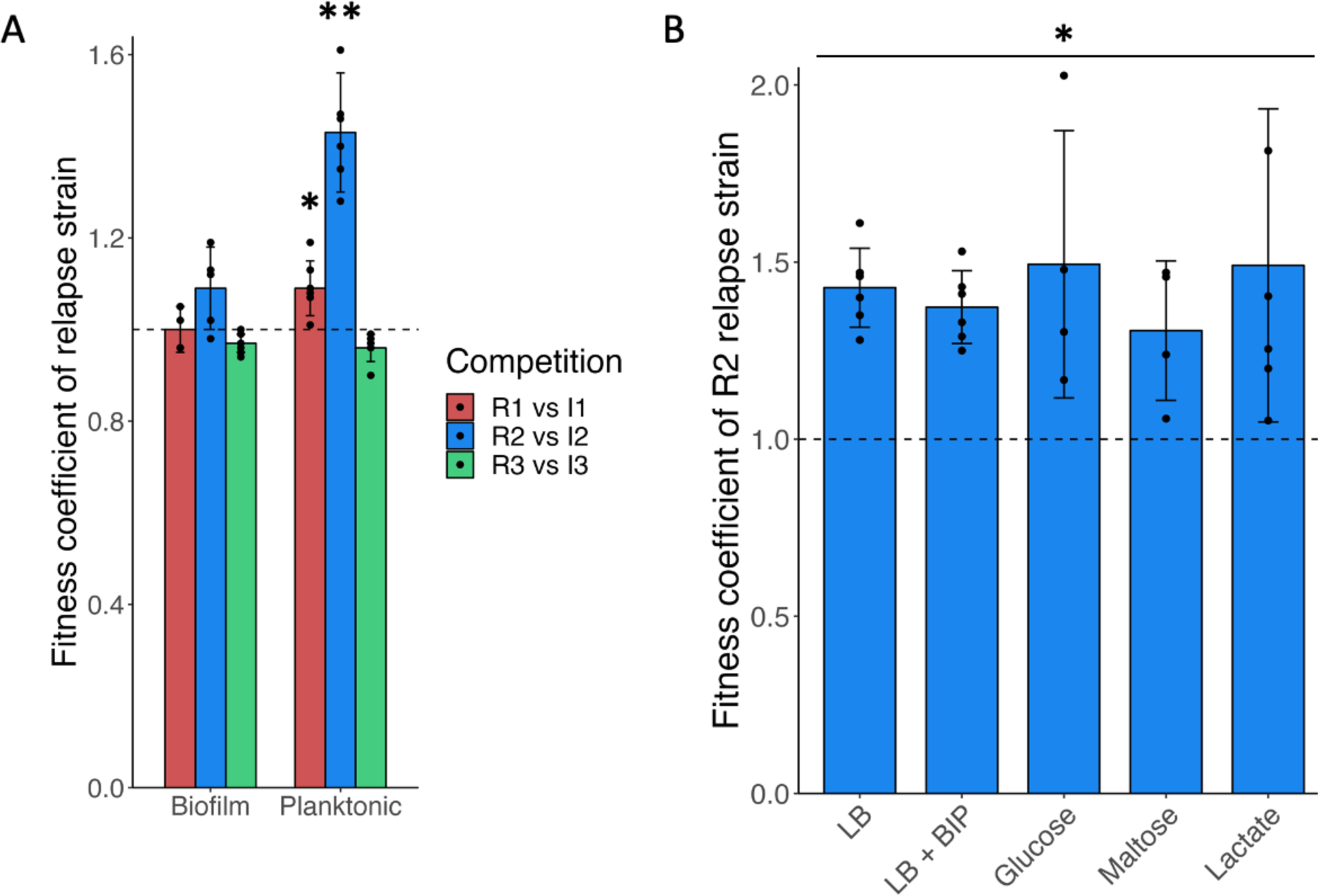
Fitness competitions and growth capacities. (**A)** Fitness competitions of relapse against ancestral strain for all three pairs in planktonic and biofilm conditions. Values are expressed as the mean of six biological replicates +/− SD with statistical significance according to Mann-Whitney test (*: p-value < 0.05, **: p-value < 0.01). **B**. Fitness competitions of R2 relapse strain against I2 ancestral strain in different conditions (same as in supplementary Figure S6.B-F). BIP: 2,2’-Bipyridyl. Values are the mean of four to six biological replicates +/− SD with statistical significance according to Mann-Whitney test (*: p-value < 0.05).

To further investigate this phenotype, we competed I2 and R2 strains in presence of different sugars as well as of an iron chelator (2,2’-Bipyridyl) based on identified genetic modifications and especially gene losses related to uptake and metabolism of these substrates (Supplementary Data 2). Surprisingly, despite these gene losses, R2 strain showed no growth defect and even globally had an advantage through a capacity to better sustain growth during the stationary phase (Supplementary Figure 6), as confirmed by competition assays (Figure 4.B). Therefore, extensive gene loss seems to have resulted in a global growth optimization, which could have helped the strain to outcompete other bacteria including its ancestor.

Another strategy to outcompete its kin and promote subsequent adaptation was identified in the relapse strain R1, which acquired a colicin-producing plasmid allowing the production of colicin E1. An overlay assay indeed confirmed the acquired capacity of R1 relapse strain to inhibit I1 strain growth (Supplementary Figure 7).

## Discussion

Treatment failure is a common issue of BJIs, allowing bacteria to remain and adapt to the host, ultimately increasing the risk of relapse and the difficulty to cure the patient. A better understanding of the mechanisms underlying pathogens’ adaptation to the host environment would allow for improving the management of relapsing BJIs. Here we investigated three cases of BJIs relapse involving pathogenic *E. coli*.

Both our genetic and phenotypic analysis highlighted a diversity of functions emerging from bacterial adaptation to the host environment during BJIs. We found that adapted strains isolated from relapses displayed a relatively high mutation rate, which is a known adaptive advantage increasing the probability to produce advantageous mutations that could be subsequently selected (13, 48, 49). However, we did not identify any mutation in genes known to be involved in DNA damage repair, suggesting that these high mutation rates were not acquired during adaptation to the host environment but rather intrinsic to these strains. This, in addition to stress-induced mutagenesis, could have further promoted the selection of beneficial traits, probably partly explaining their adaptive success.

Biofilms are known to be frequently involved in infections and relapses, including BJIs. However, none of the three investigated cases showed an increased biofilm formation capacity that could have played a role in the relapses and all strains were even characterized as poor *in vitro* biofilm formers. However, these infections were bone- or tissue-related and strong *in vitro* biofilm capacity may not necessarily translate into a selective advantage in this type of infections as compared to prosthesis-related infections, which were shown to be caused by a majority of strong biofilm formers (50). In these cases, the capacity to form biofilm correlate with the severity of the clinical outcome (51–53), while it was not the case for strains isolated from non-orthopedic biofilm infections (54, 55).

Another critical selection pressure faced by pathogenic bacteria is the host immune system. The inhibition of the production of flagellin, a known target of immune defenses, at 37°C by I1 and R1 strains, could minimize their recognition in the host environment (56, 57). This could not only contribute to immune system evasion but also to save energy through a down-regulation or suppression of flagellin production, as shown to be the case in the context of gut infection by *E. coli* (58, 59).

This immune system evasion strategy is even clearer in the case of the R2 relapse strain, which adapted and showed increased survival to human serum through the loss of two adhesins. The fact that the loss of two virulence factors could confer a selective advantage may suggest that, in these specific cases, virulence could be less important than evading the immune system as shown to happen during *in vivo* adaptation (60, 61), allowing pathogens to stay longer in the host. This trend was illustrated by the infection experiments performed in zebrafish, where no difference was observed at the exception of the R2 strain that induced a slightly higher mortality at 24hpi.

In addition, the R2 strain presented an interesting case of gene loss. Whereas genome reduction is usually considered as a strategy by which selection reduces energy consumption via the loss of genes and functions that are not used or under selective pressure (62, 63). In the case of the R2 strain, numerous gene losses seem to result in a global optimization of growth capacities, potentially correlated to lower the cost associated with the production of unnecessary genes and lower amount of DNA to replicate. It is possible that some nutritional resources could be directly acquired from the new host environment instead of being synthesized by the cells and that the loss of genes involved in this production optimizes the cell metabolism (64). Gain of growth rate or capacity in the context of an infection could be of paramount importance to outcompete other endogenous or pathogenic bacteria and thrive in the host environment. Such cases of genome reduction-induced complexification are more and more documented (20, 21) showing that innovation can also rise from gene loss and not only from gene gains or duplications followed by neo- or sub-functionalization (65).

Finally, it is an important parameter to stress the polymicrobial context of some of these BJIs infections, in which interactions between the different bacterial species or strains could play an important role in the relapses as well as in the mechanisms shaping bacterial adaptation to the host environment (66, 67). Some key phenomenon involved in the relapse or strain adaptation could therefore be dependent on bacteria-bacteria direct or indirect interactions and consequently missed in this study. However, both the limited number of strains investigated and the fact that all three cases corresponded to different BJIs could also account for the observed differences in strain profile and mechanisms involved in their *in vivo* adaptation. Further investigations with a larger cohort would help determine whether these diverse profiles and adaptive strategies are due to the various types of infection investigated here or if it constitutes a more general rule of *E. coli* pathogenic strain in BJIs. It would also help identify if some adaptive strategies are preferentially selected in relapse cases.

## Supporting information

Supplementary large dataset 2

Supplementary large dataset 1

## ACKNOWLEDGEMENTS

This work was supported by the French National Research Agency (ANR), project EvolTolAB (ANR-18-CE13-0010), by the French government’s Investissement d’Avenir Program, Laboratoire d’Excellence “Integrative Biology of Emerging Infectious Diseases” (grant n°ANR-10-LABX-62-IBEID) and by the Fondation pour la Recherche Médicale (grant DEQ20180339185). S.T.-R was supported by the French National Research Agency (ANR), project EvolTolAB (ANR-18-CE13-0010).

We thank Dr. Emma Colucci-Guyon and Dr. Laurent Boucontet for critical reading of the manuscript. We are grateful to the Zebrafish Projects Hub of the Institut Pasteur and especially to Dr. Emma Colucci-Guyon and Dr. Laurent Boucontet who carried out the infection experiments in zebrafish larvae. We are grateful to Dr. Monica Rolando for kindly providing the anti-flagellin antiserum. We are grateful to Emmanuel Chanard from Cerballiance for providing strains and microbiological information.

All authors declare no competing financial interests.

## Notes

### Competing Interest Statement

The authors have declared no competing interest.

### Summary of Updates

The Supplementary Material and Data Sets 1 and 2 have been updated

