## Supplementary Information for "Analysis of *in-patient* evolution of *Escherichia coli* reveals potential links to relapse of bone and joint infections"

[c] Département de Maladies Infectieuses et Tropicales, AP-HP, Hôpital Saint-Louis, Lariboisière, F-75010 Paris, France.

[d] FHU PROTHEE

Running Head: in-patient evolution of *E. coli* in BJIs

### Supplementary Results

#### Auto-aggregation and RDAR morphotypes on congo-red plates

Once settled and in order to colonize host environment, *E. coli* has a very large arsenal of adhesion factors. Some of them are known to promote biofilm formation via homotypic self-interactions also called auto-aggregation (1, 2), which could help facing stressful conditions such as the one encountered *in vivo*. The production of other factors such as curli and cellulose are also of critical importance (3, 4) and induces a red, dry, and rough (rdar) morphotype when the strain is plated on LB agar supplemented with congo red. However, none of these phenotypes showed any difference when comparing the relapse strains to their related ancestors, all strains even having very weak auto-aggregation capacities (supplementary Figure S1.A.B). Still, inter-couple differences were identified on congo red plates: (i) the first couple had a pink, dry and rough (pdar) morphotype characterizing the production of curli only; (ii) the second couple had a smooth and white morphotype (saw) showing the absence of matrix component production; and (iii) the third couple had pink and smooth morphotype (pas) showing the production of cellulose only.

#### Temperature specific motility

Because strain motility could have an impact on the virulence and biofilm forming capacity, we tested each strain on LB plates containing low amount of agar at 30°C and 37°C. Motility is sometime more visible at lower temperature. The second and third couple strains had no motility (supplemental Figure S2). However, an interesting phenotype was observed for the first couple. Although both strains were motile, we observed some flares characteristic of the apparition of a hypermotile clone within the population and spreading faster than the others (supplementary Figure S2). Because this phenomenon was only observed at 37°C, we investigated the importance of temperature by testing all combinations between the temperature used to produce the liquid inoculum and the temperature the motility plates were incubated at. We showed that regardless of the temperature used for the liquid culture, incubating the motility plate at 30°C always showed a homogenous motile phenotype, while plates inoculated with the same samples but incubated at 37°C always showed the apparition of flares (supplementary Figure S3.A.B.C). This suggests that the process is reversible and a westernblot analysis using anti-flagellin antibodies showed that both strains only produce flagellin at 30°C, even when pre-incubated at 37°C (supplementary Figure S3.D). Therefore, it seems that some mutants arise on plates incubated at 37°C in which the capacity to swim is restored.

Because the apparition of hyper-motile mutants suggests a high mutation rate, we used a Luria-Delbruck fluctuation test to evaluate it. Resistance to Rifampicin was used in these assays and strains I1 and R1 had a mutation rate of $2.4x10^{-6}$ and $1.7x10^{-6}$ mutations per nucleotide per generation, respectively, which is by far higher than the estimated mutation rate for resistance to Rifampicin of a wild-type *E. coli* strain (~$3.3x10^{-9}$ mutations per nucleotide per generation) (5). Such a high mutation rate could be the result of a loss of function in genes responsible for DNA repair such as the mut genes. However, no such loss of function could be identified in the genome of the clinical strains.

#### Genome analysis

To confirm that in each case the relapse strain was the same as the initial infection one all of them were subjected to whole genome sequencing (WGS) and the genomic characteristics analyzed. Each case showed that both strains were indeed the same according to their phylogroup, serotype, ANI, MLST, FimH and FumC type (Table 1).

If the genome assemblies and completeness were good for all strain, the first notable difference identified between the infection strains and their related relapse strains was the number of gained and lost genes (supplementary Table S4). Some of these genes corresponded to a whole contig that matched known plasmids, therefore constituting potential gain or loss of plasmids (supplementary Data 2). R1 relapse strain gained a potential plasmid bearing Colicin E1 production genes, R2 relapse strain lost a potential plasmid containing hypothetical protein coding genes and R3 relapse strain gained a potential plasmid containing a Toxin/Antitoxin (T/A) system (HigA/HigB). However, PacBio sequencing technology would be required to rule out what proportion of the gene gains and losses was plasmid-mediated or the consequence of big chromosomic insertions or deletions.

### Supplementary Tables

**Supplementary Table 1.** Antibiotic susceptibility profiles. Resistant (R), sensitive (S) or intermediate (I) status of each strain to commonly used antibiotics assessed by disc diffusion assay and following EUCAST breakpoints recommendations.

| **Antibiotic** | **I1** | **R1** | **I2** | **R2** | **I3** | **R3** |
|---|---|---|---|---|---|---|
| Ampicillin | R | R | R | R | R | R |
| Cefoxitin | S | S | S | S | R | R |
| Meropenem | S | S | S | S | S | S |
| Ticarcillin | R | R | R | R | R | R |
| Ticarcillin + clavulanic acid | S | S | R | R | R | R |
| Cefepime | S | S | R | R | R | R |
| Ertapenem | S | S | S | S | S | S |
| Mecillinam | S | S | S | S | S | S |
| Cefotaxime | S | S | R | R | R | R |
| Amoxicillin + clavulanic acid | S | S | R | R | R | R |
| Ceftazidime | S | S | I | I | R | R |
| Piperacillin | R | R | R | R | R | R |
| Tazocillin (Piperacillin + Tazobactam) | S | S | R | R | R | R |
| Aztreonam | S | S | R | R | R | R |
| Imipenem | S | S | S | S | S | S |
| Cefalexin | R | R | S | S | S | S |
| Tobramycin | S | S | S | S | R | R |
| Amikacin | S | S | S | S | S | S |
| Norfloxacin | R | R | S | S | R | R |
| Moxifloxacin | R | R | S | S | R | R |
| Ciprofloxacin | R | R | S | S | R | R |
| Co-trimoxazole (Trimethoprim + Sulfamethoxazole) | R | R | S | S | R | R |
| Tigecycline | S | S | S | S | S | S |
| Fosfomycin | S | S | S | S | S | S |
| Nitrofurantoin | S | S | S | S | S | S |
| Cefadroxil | S | S | S | S | S | S |

**Supplementary Table 2.** List of all strains used in this study.

| Strain | Genotype, phenotype | Relevant information | Source |
|---|---|---|---|
| MG1655 | F- lambda- *ilvG rfb*-50 *rph*-1 | E. coli Genetic Stock Center GCSC#6300 | Laboratory collection |
| TG1 | F'[*traD*36 *proAB* + *lacI*q *lacZ*DM15] *supE hsd*D5 *thi* Δ(*lac-proAB*) | Strong biofilm former | Laboratory collection |
| CFT073 | *rpoS* - | Human Blood Pyeloneprhitis (ExPEC) B2 K2 | Laboratory collection |
| MG1655 KmFRT-wbbL+ | F- lambda- *ilvG rfb*-50 *rph*-1 *wbbL*+, KmR | LPS O-antigene restored | Laboratory collection |
| MG1655 ΔfliE-R | F- lambda- *ilvG rfb*-50 *rph*-1 Δ*fliE-R*::cat, CmR | Non-motile strain | Laboratory collection |
| MG1655 yeaJ/dgcJ inter | F- lambda- *ilvG rfb*-50 *rph*-1 with and intergenic mutation between *yeaJ* and *dgcJ* | Hyper motile strain | Laboratory collection |
| 55989 | tetR | rdar morphotype on congored plates | Laboratory collection |
| MG1655 ΔcsgD | F- lambda- *ilvG rfb*-50 *rph*-1 Δ*csgD*::*aadA7*, SpecR | No cellulose production | Laboratory collection |
| MG1655 Δflu | F- lambda- *ilvG rfb*-50 *rph*-1 Δ*flu*::*GB*, KmR | No auto-aggregation | Laboratory collection |
| MG1655 PcL-flu | F- lambda- *ilvG rfb*-50 *rph*-1 KmPcL-*flu*, KmR | Strong auto-aggregation | Laboratory collection |
| MG1655 lATT Zeo-mars | F- lambda- *ilvG rfb*-50 *rph*-1 lATT-*zeo-mars*, ZeoR | Source of Zeo-mars cassette at the lambda site | Laboratory collection |
| MG1655 lATT Zeo-GFP | F- lambda- *ilvG rfb*-50 *rph*-1 lATT-*zeo-gfpmut3*, ZeoR | Source of Zeo-GFP cassette at the lambda site | Laboratory collection |
| I1 | | WT infection strain from case 1 | This study |
| I1 ΔfliC | Δ*fliC*::*zeo*, ZeoR | WT with fliC gene deleted, non-motile mutant | This study |
| I1 lATT Zeo-GFP | lATT-*zeo-gfpmut3*, ZeoR | WT with gfpmut3 gene inserted together with a zeocin resistance cassette at the λatt site under λPR constitutive promoter | This study |
| I1 lATT Zeo-mars | lATT-*zeo-mars*, ZeoR | WT with mars gene inserted together with a zeocin resistance cassette at the λatt site under λPR constitutive promoter | This study |

| | | | |
|---|---|---|---|
| R1 | | WT relapse strain from case 1 | This study |
| R1 ΔfliC | Δ*fliC*::zeo, ZeoR | WT with fliC gene deleted, non-motile mutant | This study |
| R1 lATT Zeo-GFP | lATT-*zeo-gfpmut3*, ZeoR | WT with gfpmut3 gene inserted together with a zeocin resistance cassette at the λatt site under λPR constitutive promoter | This study |
| R1 lATT Zeo-mars | lATT-*zeo-mars*, ZeoR | WT with mars gene inserted together with a zeocin resistance cassette at the λatt site under λPR constitutive promoter | This study |
| I2 | | WT infection strain from case 2 | This study |
| I2 ΔyadA-like | Δ*yadA-like*::*zeo*, ZeoR | WT with yadA-like coding gene deleted | This study |
| I2 ΔafaBCDE | Δ*afaBCDE*::*zeo*, ZeoR | WT with genes afaBCDE deleted | This study |
| R2 | | WT relapse strain from case 2 | This study |
| I3 | | WT infection strain from case 3 | This study |
| I3 lATT Zeo-GFP | lATT-*zeo-gfpmut3*, ZeoR | WT with gfpmut3 gene inserted together with a zeocin resistance cassette at the λatt site under λPR constitutive promoter | This study |
| I3 lATT Zeo-mars | lATT-*zeo-mars*, ZeoR | WT with mars gene inserted together with a zeocin resistance cassette at the λatt site under λPR constitutive promoter | This study |
| R3 | | WT relapse strain from case 3 | This study |
| R3 lATT Zeo-GFP | lATT-*zeo-gfpmut3*, ZeoR | WT with gfpmut3 gene inserted together with a zeocin resistance cassette at the λatt site under λPR constitutive promoter | This study |
| R3 lATT Zeo-mars | lATT-*zeo-mars*, ZeoR | WT with mars gene inserted together with a zeocin resistance cassette at the λatt site under λPR constitutive promoter | This study |

**Supplementary Table 3**. List of all primers used in this study.

| | |
|---|---|
| **I1/R1 and I3/R3 lATT Zeo-mars and Zeo-GFP construction and verification** | |
| lambda-ATT. B1.500-3 | CCCTGATACTCACCAGGCATCAC |
| Rev_K12-ZeoXFP-lATT | GTCACGCCAAAAGCCAATGC |
| Fw_K12-ZeoXFP-lATT | GCAAGCGCCTCGATTACTGC |
| lambda-ATT.A1.500-5 | CGATGGCGATAATATTTCACC |
| **Sanger sequencing-based fitness assessment in I2/R2** | |
| Fw_butC | CAGTCGGTTATTGCTGGCT |
| Rv_butC | CACACTTTGAGGCGACATTG |
| **Sanger sequencing-based fitness assessment calibration vs FACS in I1/R1** | |
| Fw_yehT | CAACATTTGATGCTGGAGATCG |
| Rv_yehT | GCTTCATCAATTGGCTTCAGC |
| **afaBCDE deletion in I2 and verification** | |
| Fw_afaDel_size_ZeoFRT | GATTCAGCACCTCAGTCAGAC |
| Fw_Del_afaB-E_ZeoFRT | GGTTGCTATTAACATATTTATAAAAGGTTTATTTGCCTTCAGGATAAAATATGTAGGCTGGAGCTGCTTC |
| Rv_Del_afaB-E_ZeoFRT | CAAAGGGAGCATATAGCCCCCTTCTTTCATCAGGTTAATTTTCCAGAATATCCTCCTTAGTTCCTATTCC |
| Rv_afaDel_size_ZeoFRT | GCGAGGAAGATTTCTCTTGTAGG |
| **yadA-like deletion in I2 and verification** | |
| Fw_Size_yadA_Del | TGCTGCTTAAATTGAGGTCGC |
| Fw_Del_yadA_ZeoFRT | CTCCGCACTACCGTTCTGGCTGGTGAAGTAATAATACAGGAGAATAACAGATGTAGGCTGGAGCTGCTTC |
| Rv_Del_yadA_ZeoFRT | TCCCGGGGGACTCCCCCCGGGACAGATATCTAAATCCTGACCTGGAATATCCTCCTTAGTTCCTATTCCG |
| Rev_Size_yadA_Del | CGATTCCAAATATCTCTCGCAGG |
| **fliC deletion in I1/I2 and verification** | |
| Fw_fliC | GGCATGATTATCCGTTTCTGC |
| Fw_CmFRT_ForFliC | TTGGCGTTGCCGTCAGTCTCAGTTAATCAGGTTACAACGAATTTAAATGGCGCGCCTTAC |
| Rev_CmFRT_ForFliC | CCCAATACGTAATCAACGACTTGCAATATAGGATAACGAATCGCCTACCTGTGACGGAAG |
| Rev_fliC | TCCCAGCGATGAAATACTTGC |

**Supplementary Table 4.** Genome assembly characteristics and metrics. Table showing data related to sequencing data, genome assembly quality and description.

| Case N° | Strain | Read number | Genome coverage average (SD) | Genome size (base pairs) | Contig number | GenBank Assembly accession | N50 | L50 | GC% | Complete BUSCOs | CDS | Potential plasmid gain | Potential plasmid loss | CDS gained | CDS lost | Mutation per base per year |
|---|---|---|---|---|---|---|---|---|---|---|---|---|---|---|---|---|
| 1 | I1 | 5,761,693 | 336 (170) | 5,124,061 | 283 | JARFPI000000000 | 193,231 | 9 | 50.81 | 440/440 | 4,740 | n.a | | | | |
| | R1 | 7,181,866 | 418 (200) | 5,133,911 | 268 | JARFPF000000000 | 193,231 | 10 | 50.81 | 440/440 | 4,754 | 1 | 0 | 35 | 19 | $8.38x10^{-6}$ |
| 2 | I2 | 5,591,377 | 316 (144) | 5,300,036 | 457 | JARFPH000000000 | 194,756 | 9 | 50.69 | 440/440 | 4,914 | n.a | | | | |
| | R2 | 5,271,593 | 308 (132) | 5,122,701 | 304 | JARFPE000000000 | 208,913 | 8 | 50.77 | 440/440 | 4,775 | 0 | 1 | 31 | 170 | $1.11x10^{-6}$ |
| 3 | I3 | 5,594,583 | 341 (138) | 4,900,901 | 244 | JARFPG000000000 | 137,146 | 11 | 50.79 | 440/440 | 4,511 | n.a | | | | |
| | R3 | 4,748,473 | 289 (118) | 4,904,465 | 239 | JARFPD000000000 | 137,146 | 11 | 50.79 | 440/440 | 4,518 | 1 | 0 | 14 | 9 | $5.64x10^{-6}$ |

**Legends for Supplementary Data Sets**

**Supplementary Data Set 1**
**Antimicrobial resistance prediction**. Resistance status of each strain as predicted by ResFinder based on whole genome sequences. Each sheet in the file corresponds to one case with the corresponding infection and relapse strain.

**Supplementary Data Set 2**
**Genes gained and lost by each relapse strain**. Each sheet in the file corresponds to one case with the corresponding infection and relapse strain. For each case, the identifier of the genes that were gained or lost by the relapse strain are shown together with their annotation. Sets of genes corresponding to a potential plasmid are framed.

### Supplementary figures

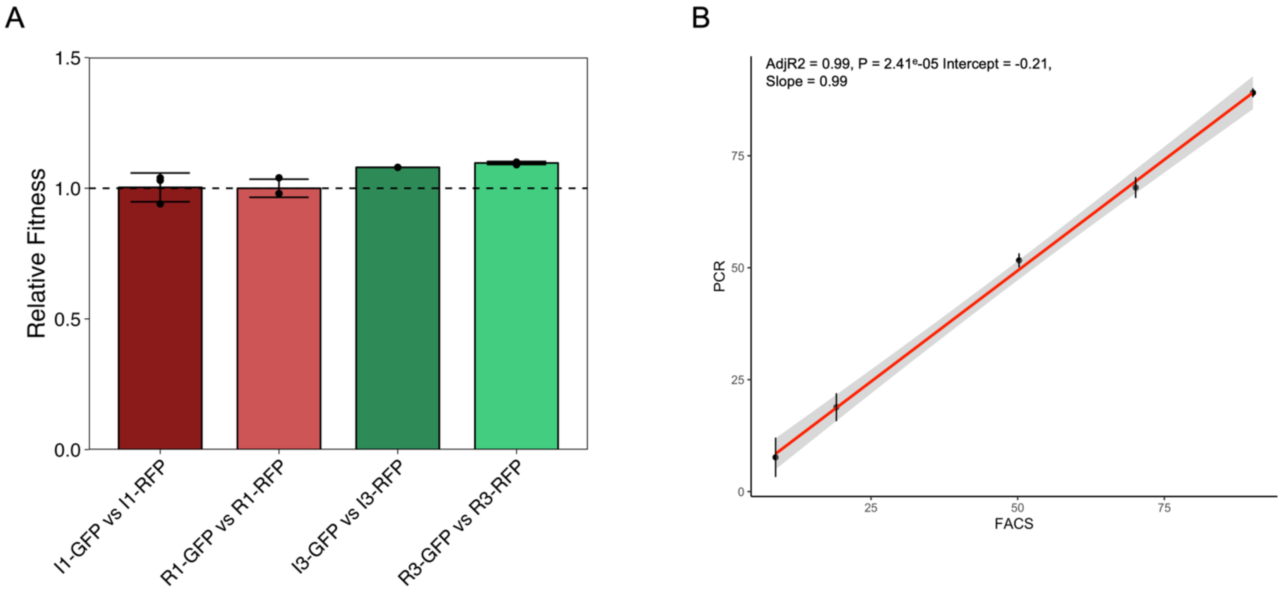

**Supplementary Figure 1**. Optimization of competition protocols. (**A**) Assessment of the fitness cost associated to the expression of each fluorescence marker (GFP or mars) in each strain. There is a slight cost to produce mars marker in I3 and R3 strains as compared to the production of GFP. Therefore, both combinations of fluorescence marker were used in the competitions assays for all strains. (**B**) Comparison of FACS approach and Sanger based QSVanalyzer approach to assess the proprotion of each strain in a set of mixed samples with different proportions of I1 and R1 strains taged with GFP and mars, respectively.

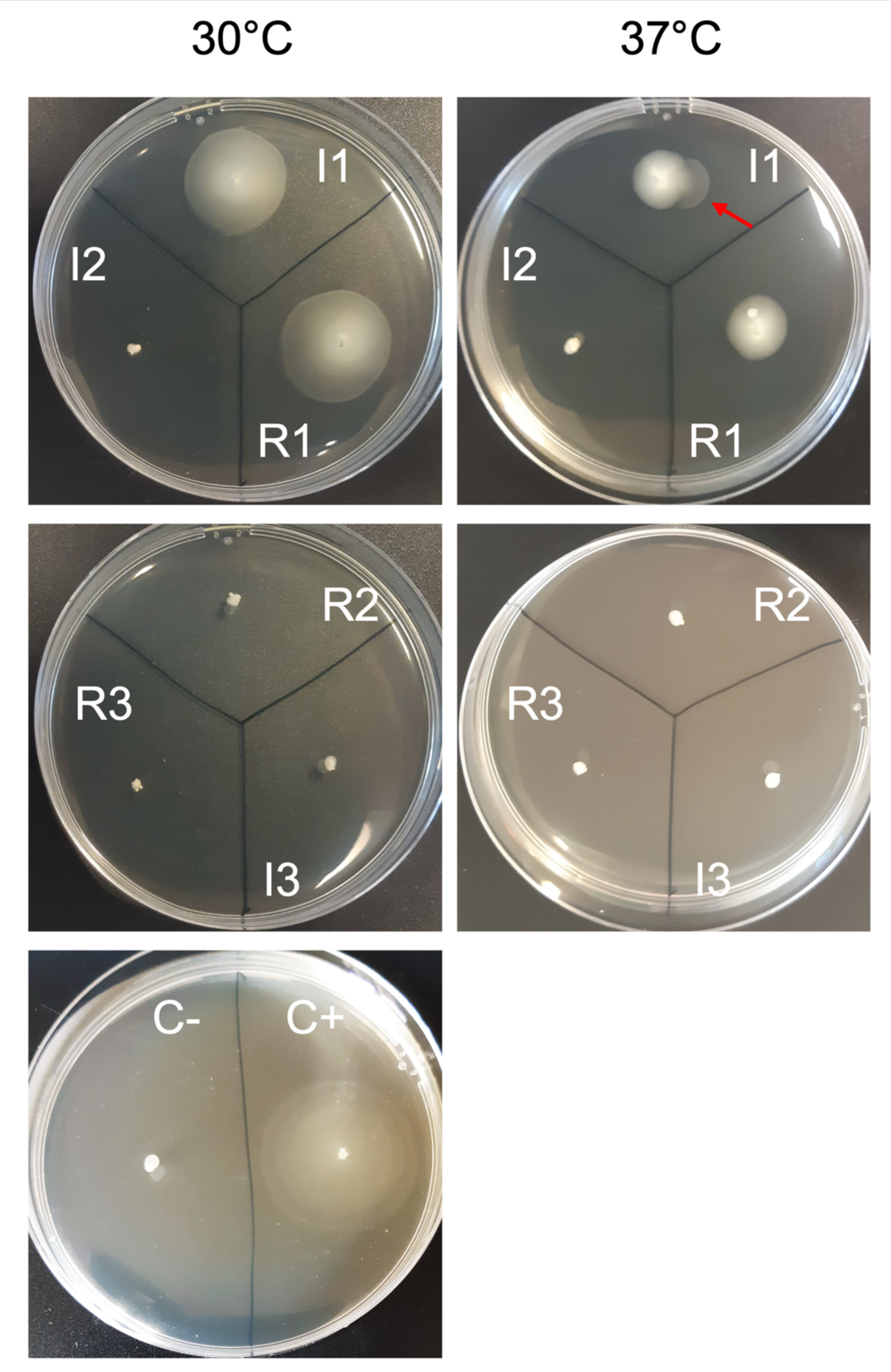

**Supplemental Figure 2**. Motility of each strain assessed on low agar plates at 30 and 37°C. A *E. coli* MG1655 strain deleted for fliE to fliR genes is used as a negative control and a *E. coli* MG1655 strain in which the motility is increased by the insertion of an IS1 in *yeaJ/dgcJ* encoding a diguanylate cyclase resulting in an increase of c-di-GMP, which in turn fuels motility was used as a positive control.

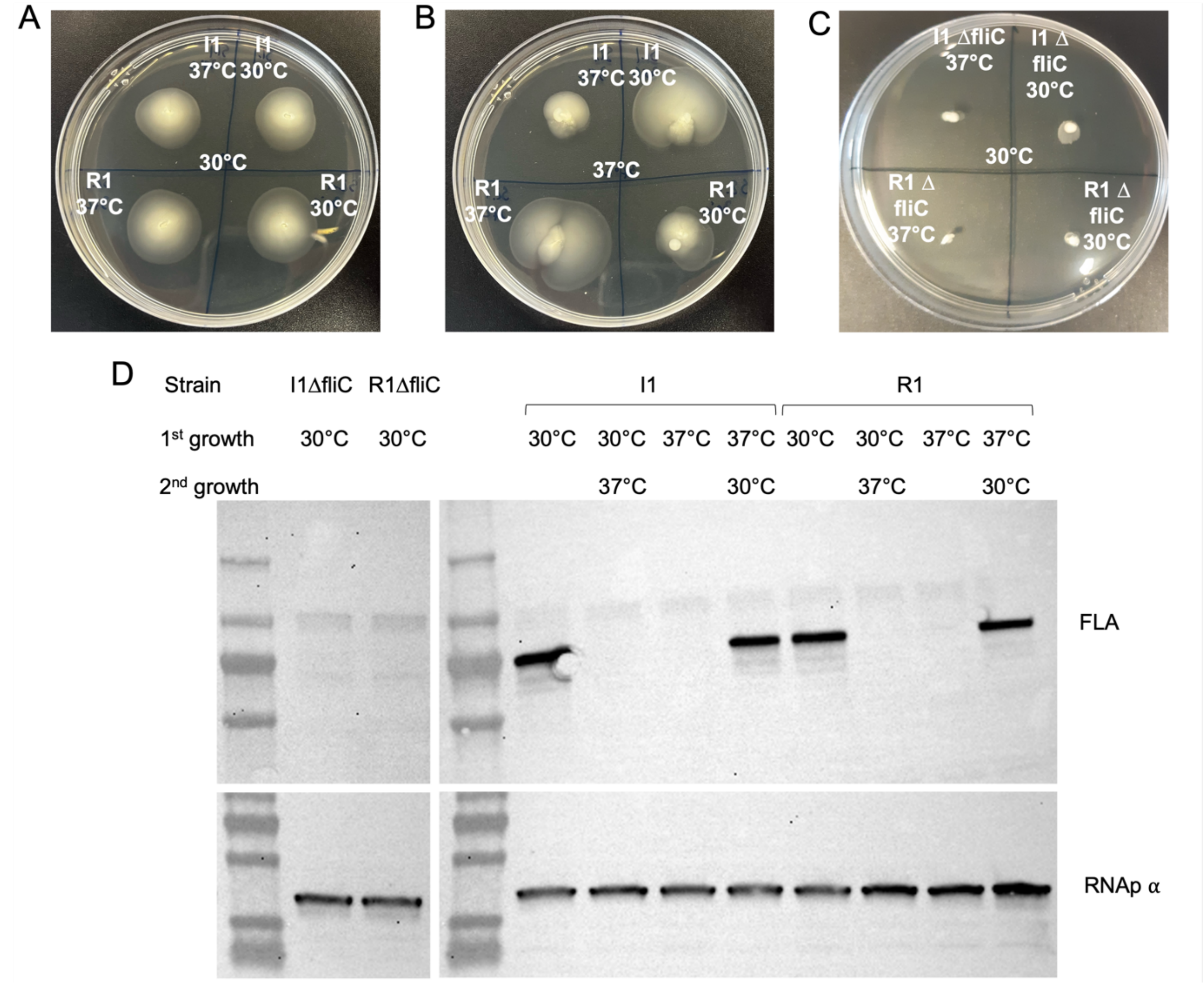

**Supplementary Figure 3**. Influence of temperature on I1 and R1 strain motility. Motility of each strain at 30°C (**A**) or 37°C (**B**) when pre-cultured in liquid at 30°C or 37°C. (**C**) Motility of each strain deleted for *fliC* when pre-cultured at 30°C or 37°C and incubated at 30°C for motility testing. (**D**) Production of flagellin (top panel) and RNA polymerase alpha subunit used as a loading control (bottom panel) in I1 and R1 strains assessed by westernblot analysis in different combinations of temperature in liquid cultures.

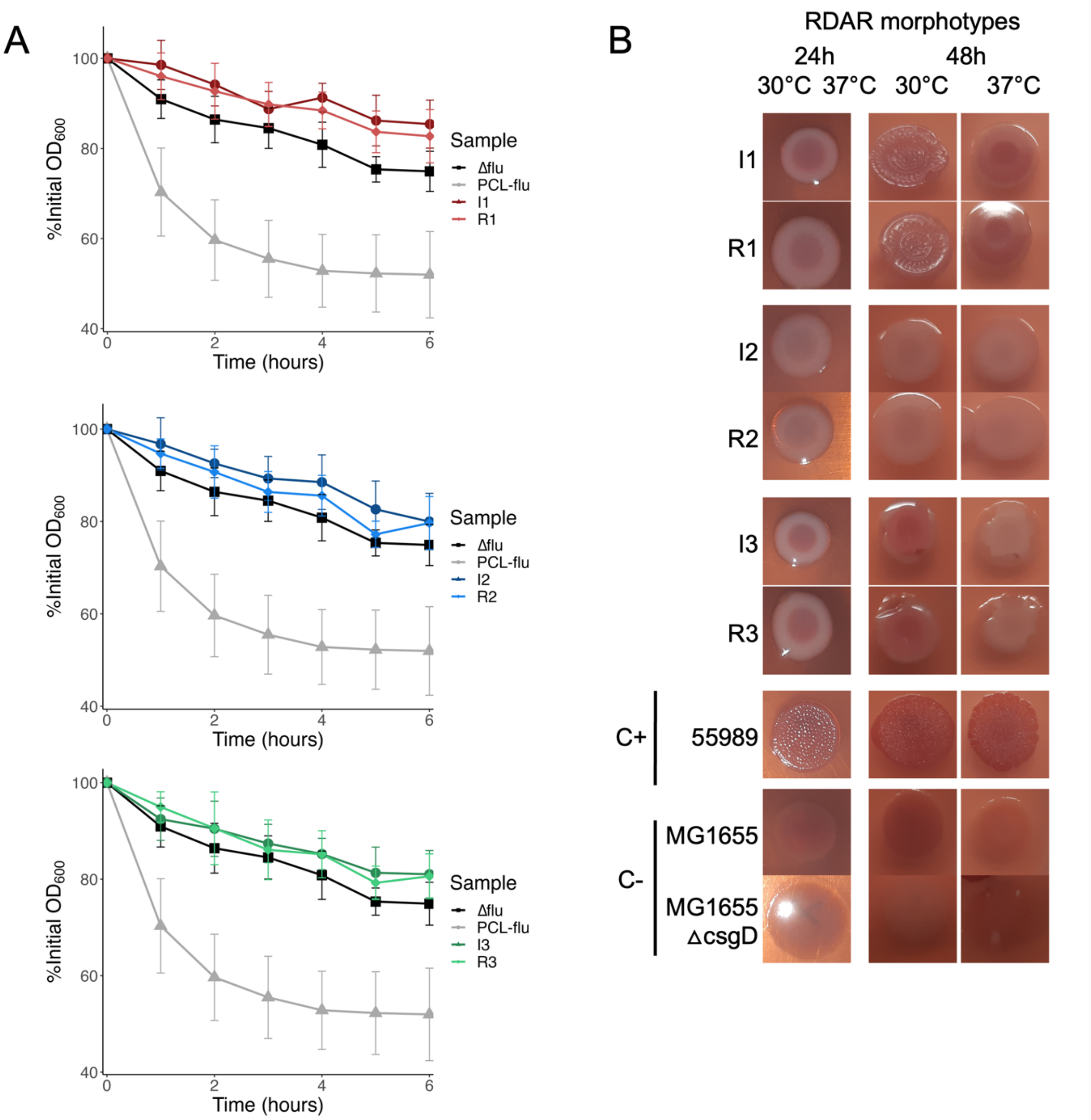

**Supplementary Figure 4**. **A**. Auto-aggregation kinetic of all clinical strains. In each case, the infection strain is compared to the relapse strain as well as a strain of *E. coli* MG1655 in which *flu* is under the control of a constitutive promoter (PCL-flu) used as a positive control and a strain of *E. coli* MG1655 in which *flu* is deleted (Δflu) used as a negative control. Each time point is the mean of 3 biological replicate +/- SD. **B**. Pictures showing the red, dry, and rough (rdar) morphotypes of each strain after 24 or 48 hours of incubation at either 30°C or 37°C. The first couple had a pink, dry and rough (pdar) morphotype characterizing the production of curli only. The second couple had a smooth and white morphotype (saw) showing the absence of matrix component production. The third couple had pink and smooth morphotype (pas) showing the production of cellulose only

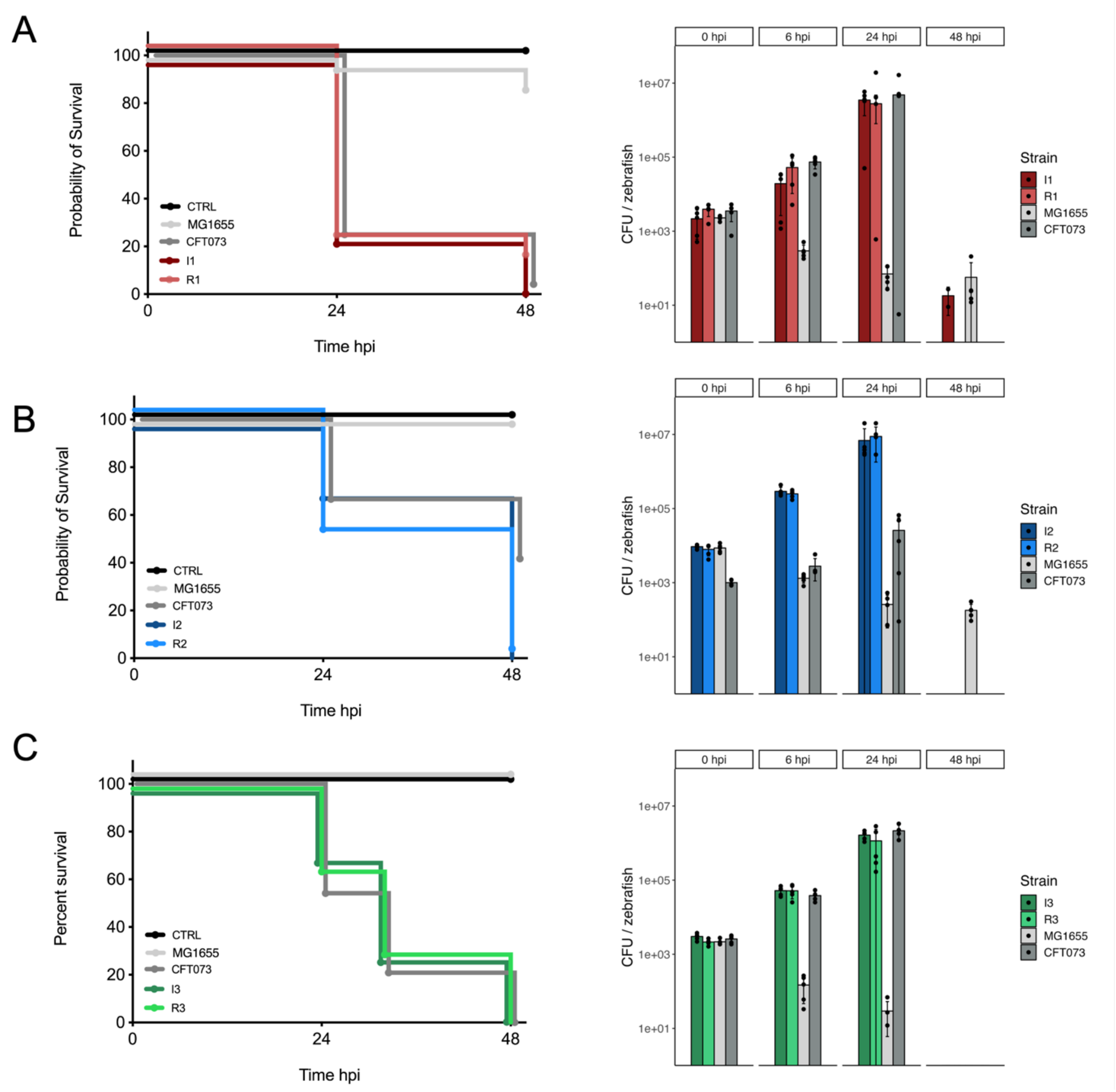

**<u>Supplementary Figure 5</u>**. Zebrafish *in vivo* experiments. Monitoring of zebrafish survival (left panel) and bacterial burden into the zebrafish (right panel) for I1 and R1 (**A**), I2 and R2 (**B**) and I3 and R3 (**C**). MG1655 strain was used as a negative control and CFT073 strain as a positive control. Fifteen zebrafish were used for each experiment.

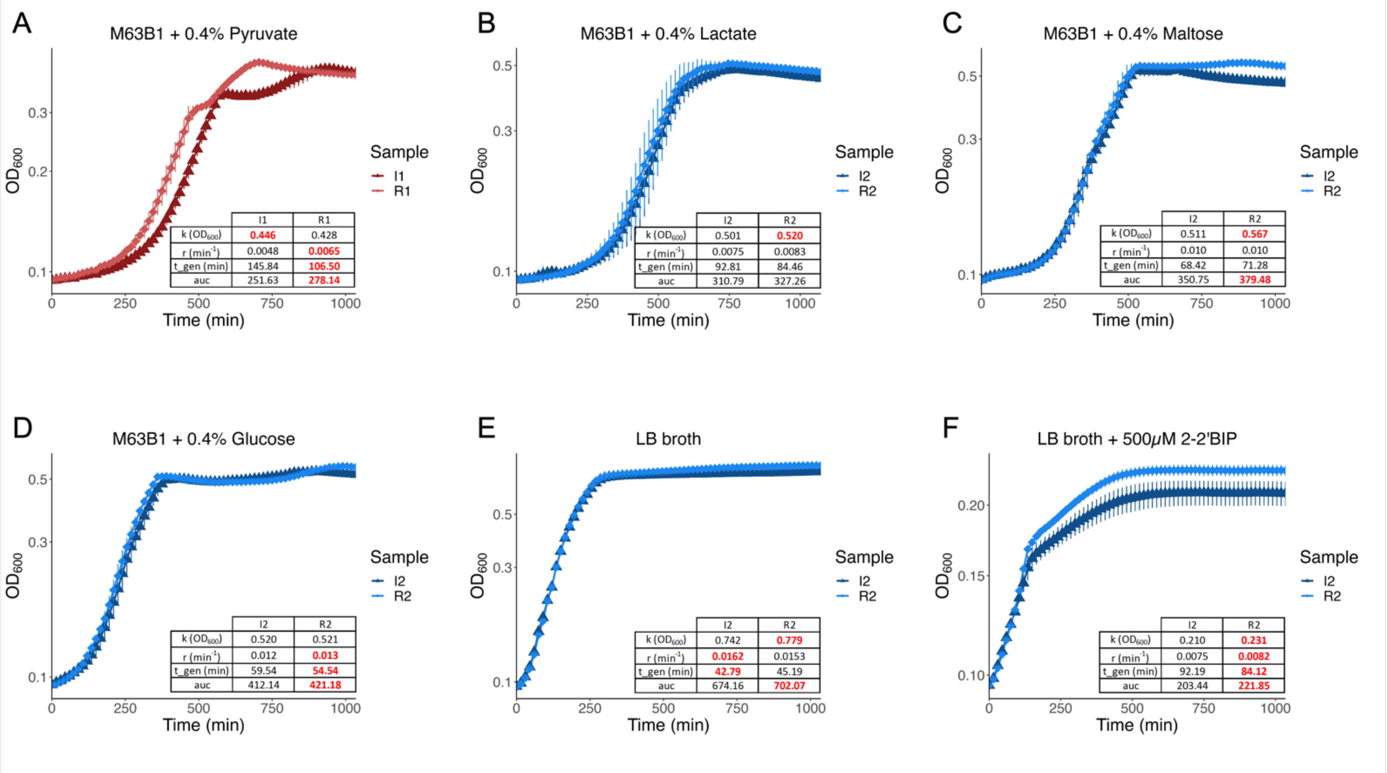

**<u>Supplementary Figure 6</u>**. Growth capacities in different conditions. (**A)** Growth curve and related parameters inferred by the R package GrowthCurver (values for which the difference between both strains was significant (Mann-Whitney test, p-value < 0.05) are highlighted in bold red) for I1 and R1 strains grown in M63B1 minimum medium supplemented with 0.4% pyruvate. Values are mean of five biological replicates +/- SD. (**B-F)** Growth curve and related parameters inferred by the R package GrowthCurver (values for which the difference between both strains was significant (Mann-Whitney test, p-value < 0.05) are highlighted in bold red) for I2 and R2 strains grown in M63B1 minimum medium supplemented with 0.4% lactate (B), Maltose (C), Glucose (D) as well as in LB medium (E) and LB medium supplemented with 500 µM of 2,2′-Bipyridyl (F). Values are the mean of four to five biological replicates +/- SD.

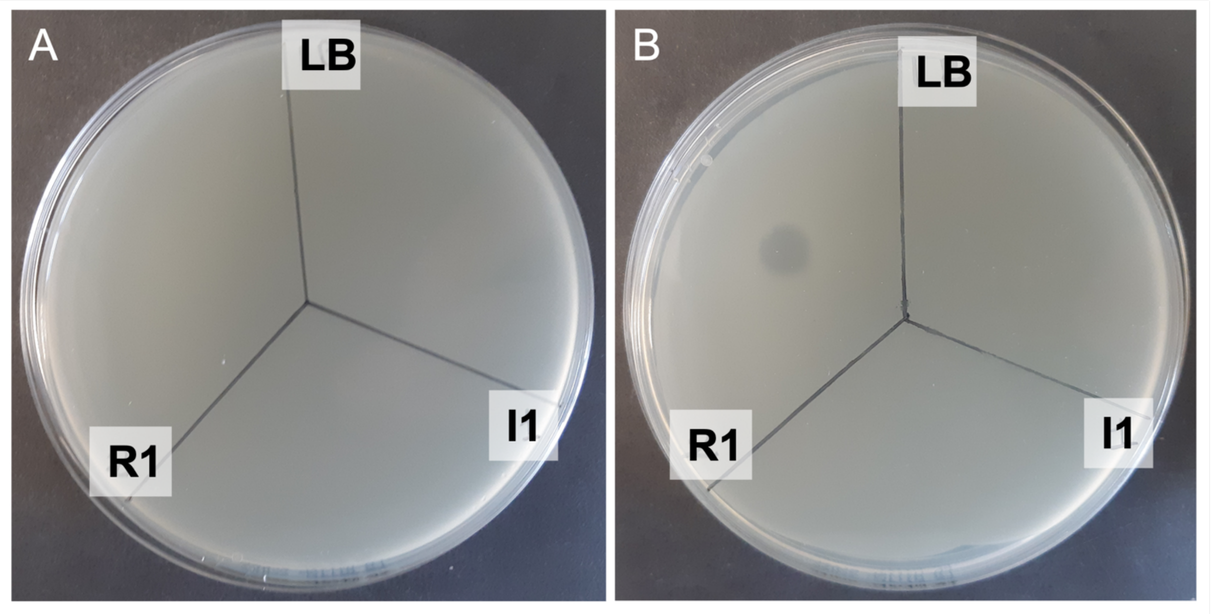

**Supplemental Figure 7**. Colicin production in R1 relapse strain. **A**. Overlay of R1 strain showing no growth inhibition by a drop of either I1 or R1 supernatant from a culture in presence of Mitomycin C to induce colicin production. **B**. Same experiment with an overlay of I1 strain showing a growth inhibition by a drop of R1 supernatant from a culture in presence of Mitomycin C to induce colicin production. In both cases, LB is used as a negative control.
